# Structure of *Mycobacterium tuberculosis* Cya, an evolutionary ancestor of the mammalian membrane adenylyl cyclases

**DOI:** 10.1101/2021.12.01.470738

**Authors:** Ved Mehta, Basavraj Khanppnavar, Dina Schuster, Ilayda Kantarci, Irene Vercellino, Angela Kosturanova, Tarun Iype, Sasa Stefanic, Paola Picotti, Volodymyr M. Korkhov

## Abstract

*Mycobacterium tuberculosis* adenylyl cyclase (AC) Rv1625c / Cya is an evolutionary ancestor of the mammalian membrane ACs and a model system for studies of their structure and function. Although the vital role of ACs in cellular signaling is well established, the function of their transmembrane (TM) regions remains unknown. Here we describe the cryo-EM structure of Cya bound to a stabilizing nanobody at 3.6 Å resolution. The TM helices 1-5 form a structurally conserved domain that facilitates the assembly of the helical and catalytic domains. The TM region contains discrete pockets accessible from the extracellular and cytosolic side of the membrane. Neutralization of the negatively charged extracellular pocket Ex1 destabilizes the cytosolic helical domain and reduces the catalytic activity of the enzyme. The TM domain acts as a functional component of Cya, guiding the assembly of the catalytic domain and providing the means for direct regulation of catalytic activity in response to extracellular ligands.

**One-Sentence Summary:** Structure of *M. tuberculosis* membrane adenylyl cyclase Cya

---

Adenylyl cyclases (ACs) convert molecules of ATP into 3,5-cyclic AMP (cAMP), a universal second messenger and a master regulator of cellular homeostasis (*1*). In mammalian cells, the membrane-associated ACs (Fig. 1A) generate cAMP upon activation of the cell surface receptors, GPCR, via G protein subunits (*2*), or in some cases by Ca^2+^/calmodulin (*3*). The cAMP molecules produced by the ACs bind to a number of effector proteins, including protein kinase A (*4*), cyclic nucleotide-gated channel channels (*5*), EPAC (*6*), popeye proteins (*7*) among others, which in turn regulate virtually all aspects of cellular physiology (*8*). The nine mammalian membrane ACs (AC1-9) share the topology and domain organization: twelve transmembrane (TM) helices with TM6 and TM12 extending to form a coiled coil of the helical domain (HD), linking the TM bundle to the bipartite catalytic domain (Fig. 1A) (*9*). Recently we determined the cryo-EM structure of the full-length AC, the bovine AC9 bound to G protein as subunit, revealing the organization of the membrane-integral region of a membrane AC (*9*). Although the structure provided important insights into the mechanism of AC9 auto-regulation, the role of the elaborate twelve-helical membrane domain remains unexplained.

**Fig. 1.**
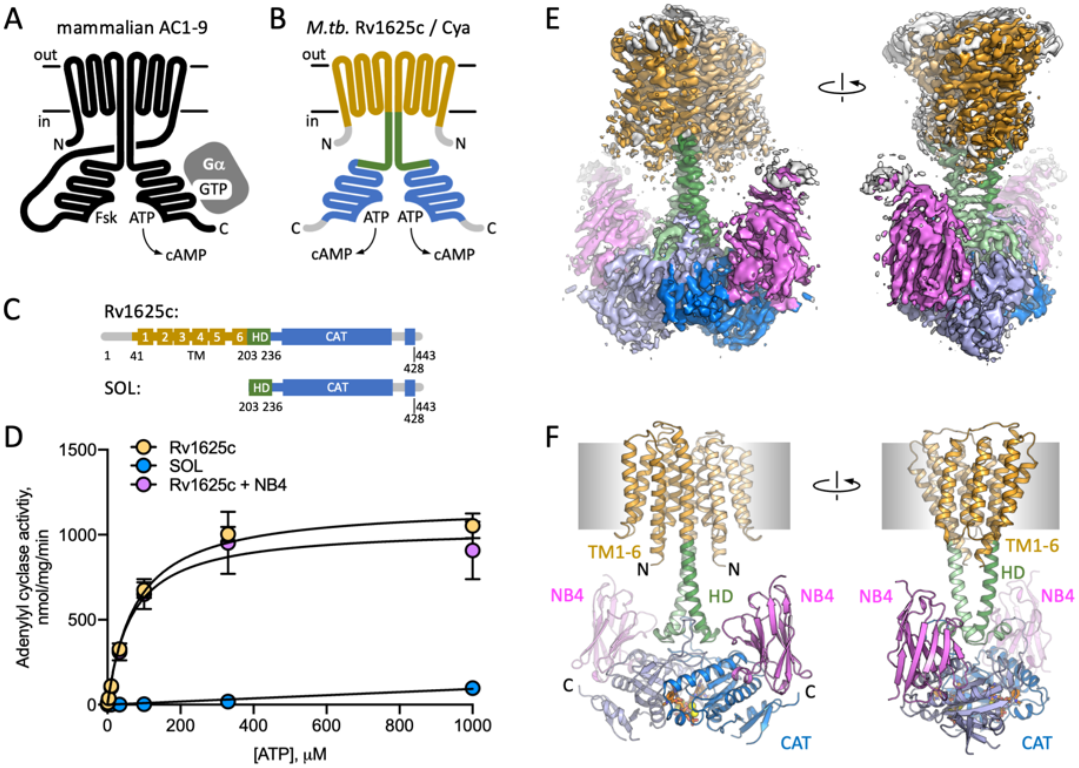
Structure of Cya-NB4 complex. (**A**) Schematic representation of the mammalian membrane ACs, indicating the key elements of AC structure: 12 TM domains, two catalytic domains, an ATP and a forskolin (Fsk) binding site. The protein is depicted in a G protein-bound state. (**B-C**) A schematic representation of Rv1625c / Cya, illustrating the regions resolved in the cryo-EM structure. The TM region is coloured orange, the helical domain (HD) is green, the catalytic domain is blue. Regions absent in the cryo-EM structure are grey. (**D**) The activity of the full-length Cya in detergent is similar in the absence (yellow) and in the presence of nanobody NB4 (pink); the soluble domain of Cya (SOL, blue) shows low levels of activity. For all experiments the data are shown as mean ± S.E.M. (n = 3). (**E**) The density map of Cya-NB4 complex at 3.57 Å resolution, obtained using masked refinement of the best dataset with c2 symmetry imposed. (**F**) The corresponding views of the atomic model of Cya-NB4 complex, coloured as in B-C. “N” indicates the N-terminal part of the protein; “HD” – helical domain; “CAT” – catalytic domain.

A putative evolutionary ancestor of the mammalian membrane ACs has been identified in the genome of *Mycobacterium tuberculosis*: Rv1625c, or Cya (*10*); for simplicity, we refer to this protein as Cya throughout. This protein is one of the sixteen ACs present in genome of *M. tuberculosis*, and one of five ACs predicted to be polytopic membrane proteins (*11*). The exact function of Cya is not clear, although available evidence indicates that the protein may be involved in CO_2_ sensing (*12*) and cholesterol utilization by *M. tuberculosis* (*13, 14*). Cholesterol utilization during infection by *M. tuberculosis* is linked to its pathogenesis (*15*), indicating a potential role of Cya at some stages of macrophage infection. The catalytic domain of Cya belongs to the same fold as those of the mammalian ACs, the class III AC / guanylyl cyclase (GC). Unlike the mammalian ACs, Cya is predicted to include only six TM helices, with TM6 extending into a HD connected to the catalytic domain (Fig. 1B-C). The protein has to dimerize to form a functional unit that has been previously presumed to resemble the pseudo-heterodimeric fold of the full-length mammalian ACs (*16, 17*). Although the structures of the *M. tuberculosis* Cya soluble domain in monomeric form (*16*) and that of the homologous *M. intracellulare* Cya in dimeric form (*17*) have been solved using X-ray crystallography, a structure of a full-length mycobacterial AC that includes the TM region has not been determined until now.

The role of the membrane domain in membrane ACs is a mystery. Polytopic membrane nucleotidyl cyclases with membrane domains of known function have been described in several organisms. The ACs in *Paramecium, Plasmodium* and *Tetrahymena* are fused to an ion channel module (*18*), and the light-sensitive GCs in several fungi, such as *Blastocladiella emersonii*, which are fused to a rhodopsin-like membrane module that binds a light-sensitive retinal chromophore (*19*). However, the functional role of the TM regions in the mammalian membrane ACs remains unclear. A recent structure of the bovine AC9 shed relatively little light on the possible function of the membrane domain (*9*), but provided a description of the unique TM helix arrangement in the membrane anchor of the protein. Interestingly, experiments with domain-substituted Cya, a presumptive evolutionary ancestor of the mammalian ACs, have suggested that its membrane region may have a regulatory role, potentially acting as a receptor for yet unidentified ligands (*20*).

Understanding the structure and function of the AC membrane domains, conserved through evolution from bacteria to mammals, is essential for understanding the regulation of cAMP generation by the cells at rest and during AC activation. The importance and necessity of a complex polytopic membrane domain in the membrane ACs is one of the key open questions in the cAMP signaling field. To address this key question, we set out to determine the structure of the model membrane AC, *M. tuberculosis* Cya.

## Results

### Characterization of the full-length Cya

The full-length *M. tuberculosis* Cya (Fig. 1C) was expressed in *Escherichia coli* and purified in digitonin micelles using affinity chromatography followed by size exclusion chromatography (Fig. S1). Adenylyl cyclase activity assays confirmed that the full-length protein was purified in a functional form (Fig. 1D; K_m_ for Cya was ∼80 μM). The “SOL” construct, consisting of the catalytic cytosolic domain (residues 203-428) showed low activity (Fig. 1D), indicating the importance of the membrane-spanning region for proper assembly and activity of the cyclase, in agreement with the previous reports (*17, 21*).

### Nanobody NB4 facilitates Cya structure determination

Cya is a relatively small membrane protein (45 kDa for a monomer). The presence of a helical domain linking the TM domain with the catalytic domain of Cya makes this protein a challenging target for structural studies. To increase the likelihood of high resolution structure determination, we used the purified Cya to generate a panel of nanobodies, camelid antibody fragments (*22*), recognizing the target protein with high affinity. One of these reagents, nanobody 4 (NB4), had no effect on the catalytic activity of the full-length cyclase (Fig. 1D), but had a nanomolar affinity for the SOL domain (Fig. S1E). We reconstituted a complex of the detergent-purified full-length Cya and NB4 (mixed at a molar ratio of 1:1.5), in the presence of 0.5 mM MANT-GTP (a non-cyclizable nucleotide-derived AC inhibitor), and 5 mM MnCl_2_. The sample was subjected to cryo-EM imaging and single particle analysis (Fig. S2), yielding a 3D reconstruction of the protein in C1 symmetry at 3.8 Å resolution (Fig. S2-3, Table S1).

The reconstruction revealed the full-length Cya arranged as a dimer bound to three copies of NB4 nanobody: two copies bound symmetrically to the SOL portions of the Cya dimer, and one asymmetrically bound to the extracellular surface of the protein (Fig. S3). To visualize the details of the Cya-NB4 interaction, we crystallized the SOL construct in the presence of NB4 and solved the X-ray structure of the complex at 2.1 Å resolution (Fig. S4A, Table S2). The structure showed an extensive interaction interface between the negatively charged surface of the monomeric SOL domain and the NB4 (Fig. S4B). Interestingly, the crystallized construct did not form a native-like dimeric form of the enzyme, but nevertheless retained the ability to bind to MANT-GTP/Mn^2+^, with an unusual twist of the MANT-GTP base (Fig. S4C). The well-resolved structure of the Cya SOL-NB4 complex allowed us to reliably place NB4 into the cryo-EM density map (Fig. S3-4).

To improve the resolution of the cryo-EM density map in the regions of highest interest, we masked out the extracellular NB4 density and refined the Cya-NB4 dataset imposing the C2 symmetry. This resulted in a 3D reconstruction at 3.57 Å resolution, which allowed us to reliably trace the polypeptide chain in the cryo-EM density map, covering residues 41-428 of the full-length Cya construct (Fig. 1E-F, Fig. S3, Table S1).

### Key features of the Cya structure

Our 3D reconstruction revealed the previously unresolved portion of the protein, the 6-TM bundle, arranged into a homodimer (Fig. 1F, 2B, 2D). The SOL portion of the protein, linked to the TM region via the helical domain (HD), adopted a conformation consistent with our previous structure of the SOL domain of *M. intracellulare* Cya homologue (Fig. S4) (*17*). The two nucleotide binding sites of Cya are occupied with the molecules of MANT-GTP/Mn^2+^, which we modelled based on the previous structures and the X-ray structure of SOL-NB4 complex (Fig. S4). The C1 and the C2 maps provide no clear evidence of asymmetry in the active site, which we have observed in the structure of *M. intracellulare* Cya (*17*). Therefore, the two MANT-GTP molecules were modelled in identical orientations.

The HD region is believed to be a critical element in the membrane and soluble ACs and GCs, as this region couples the N-terminal regulatory domains to the catalytic function of these proteins (*23-25*). In Cya, the HD extends from the TM6 (Fig. 2A-B), forming a coiled-coil observed in the structures of homologous proteins, including AC9 (Fig. 2C) (*9*) and sGC (*23*) (Fig. 3). Interestingly, the size difference between the HD helix in Cya and the HD1 and HD2 helices in AC9 leads to an ∼90 rotation of the corresponding TM regions, relative to the catalytic domains (Fig. S5A). This may be an indication that the exact structural alignment of the TM domain and the relatively remote catalytic domain may not be a conserved feature of the membrane ACs. Instead, it is likely that the precise TM-HD and HD-catalytic domain coupling plays the key role in the formation and regulation of the catalytic center in the membrane AC, consistent with the function of the HD as a transducer element in the AC structure (*26*).

**Fig. 2.**
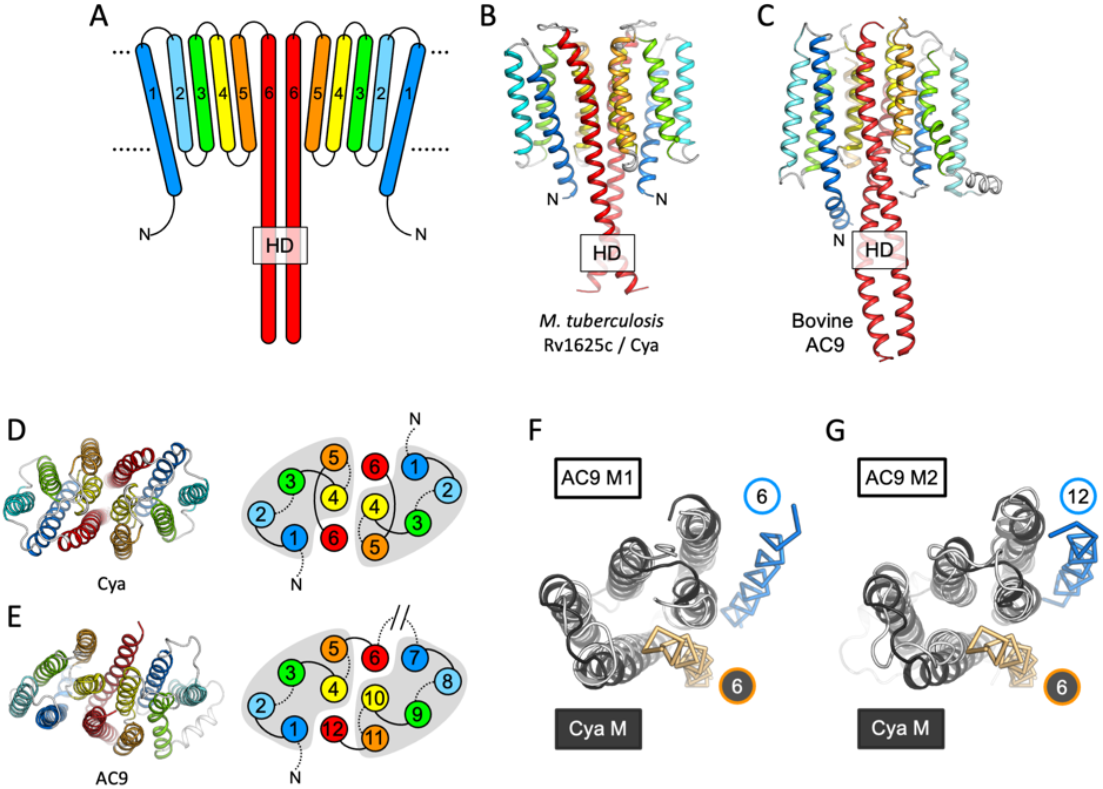
Features of the Cya TM domain. (**A**) A schematic representation of the 6-TM bundles (TM1-TM6) of Cya, arranged as dimer. (**B-C**) A view of Cya (B) and AC9 (C) TM domain parallel to the membrane plane. (**D-E**) A view of Cya (D) and AC9 (E) TM domain perpendicular to the membrane plane. The schematic indicates the relative arrangement of the TM helices, with helices 4, 5 and 6 at the dimer interface. The grey shapes indicate the conserved structural motif (TM1-5 in Cya, TM1-5 and TM7-11 in AC9) of the membrane ACs. The extracellular and intracellular loops connecting the TM helices are shown using solid and dotted lines, respectively. The connection between TM6 and TM7 of AC9 is indicated as a broken line, in place of the catalytic domain C1a and the connecting loop C1b. (**F-G**) Alignment of the 6-TM bundles of Cya (black) and AC9 (white) reveals a high level of structural conservation, in particular in the TM1-5 (RMSD 3.42 Å over 112 residues; F) and TM7-11 regions of the two proteins (RMSD 3.56 over 112 residues; G). The positions of the HD-forming TM helices are conserved, but the TM6 and TM12 are swapped in AC9 (blue) relative to Cya (orange).

**Fig. 3.**
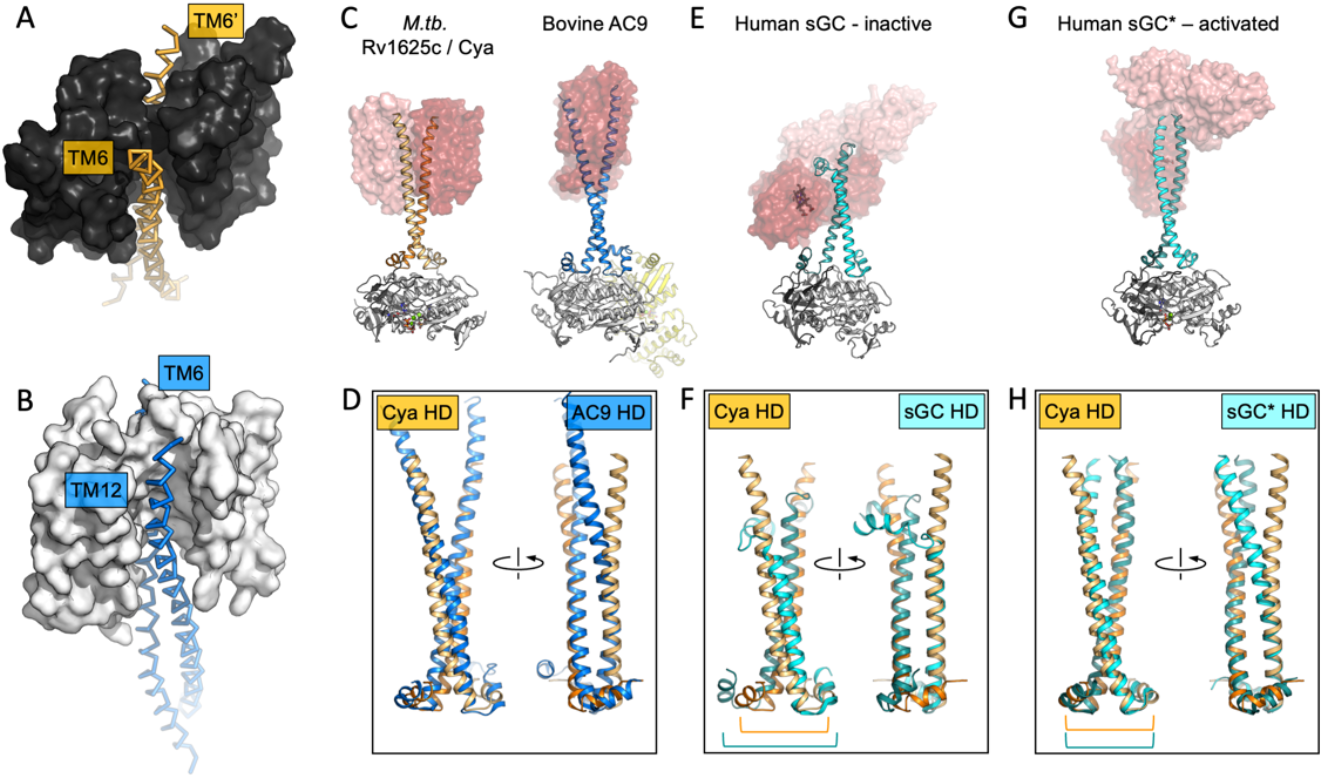
The role of the TM domain as a chaperone for HD / AC assembly. (**A-B**) Guide rail-like structures are formed by the TM1-5 in Cya (black, A) and the TM1-5/TM7-11 in AC9 (white, B). The arrangement of these rail-like structures positions the TM6 helices for optimal assembly of the helical (HD) and the catalytic domain. (**C**) The views of Cya and AC9 with the TM1-5 (Cya) and TM1-5/TM7-11 (AC9; PDB ID: 6r3q) represented as transparent surface. (**D**) The TM-HD regions of Cya and AC9 aligned. Despite the difference in HD length and the deviation in the TM domains, the cores of the HD domains are well aligned. (**E-F**) Similar to C-D, for the human soluble guanylyl cyclase sGC (inactive form; PDB ID: 6jt0). (**G-H**) Similar to E-F, for the activated form of sGC (sGC*; PDB ID: 6jt2). The brackets in F and H indicate the misaligned (F) and aligned (H) portions of the Cya and sGC HDs.

The resolved portion of the Cya N-terminus (residues V_41_ARRQR_46_), rich in positively charged residues, is immediately adjacent to the HD region. The early work on Cya identified the mutations in this region that disrupt the function of the protein (*17*), suggesting that the intact residues in the N-terminus stabilize the HD. Our structure provides the structural basis for understanding the likely disruptive effects of these mutations. The positively charged residues R43-R44 likely stabilize the negatively charged surface of the HD (Fig. S5B).

### The TM1-5 bundle as a rail for the HD helices

The TM helices 4, 5 and 6 of Cya form an extensive dimer interface within the membrane (Fig. 2D). The dimer interface residues, close to the “core” of the protein, are relatively well conserved among the Cya homologues from Mycobacteria (Fig. S7), with relatively poorly conserved residues in TM1-3. A comparison of the 6-TM bundle of Cya with the corresponding regions in the bovine AC9 (TM1-6 and TM7-12) shows that the helices TM1-5 (and TM7-11 for the AC9) form a well defined structural motif (Fig. 2D-E). A striking difference between the Cya and AC9 membrane domains is that the TM region that forms the HD helix is swapped in AC9: the TM12 of AC9 occupies the same position as the TM6 of Cya. Similarly, TM6 in AC9 is placed in a corresponding position relative to the TM7-11 (Fig. 2F-G). The TM1-5 bundle in Cya appears to act as a “guide rail” for the TM6/HD helix of Cya, guiding the correct assembly of the HD coiled coil and the catalytic domain of the cyclase (Fig. 3A). This feature is remarkably similar in AC9, with TM1-5 and TM7-11 arranged in a near-identical way (Fig. 3B), and with a closely matching HD core (Fig. 3D).

The previous experiments in *M. intracellulare* Cya have shown that the HD and the TM regions of the protein are critically important for the protein’s dimerization and functional assembly (*17*). The lack of the TM region results in failure to form a stable active dimer of *M. tuberculosis* Rv1625c / Cya, even in the presence of a nucleotide analogue MANT-GTP, judged by the inability of MANT-GTP to induce crystallization of the protein in a dimeric form (Fig. S4). In contrast, the soluble domain of the *M. intracellulare* Cya is effectively dimerized by MANT-GTP (*17*). The importance of the TM domain as a factor that promotes correct protein folding is further illustrated by the ability of the isolated Cya SOL construct to form an inactive domain-swapped dimeric assembly (*27*). It is thus tempting to suggest that the key function of the TM domain in a membrane AC is to guide the assembly of the enzyme in a catalytically competent form.

This may have important implications for AC regulation. In a related enzyme, the NO-sensing sGC, the heme-containing NO-receptor domain is fused to the HD region in place of the TM regions seen in Cya or in the mammalian AC9 (Fig. 3E-G). In its inactive form, the sGC displays a conformation where HD helices are bent, with an accompanying substantial unwinding of the helical domain core (Fig. 3E). Comparison of the Cya HD core with that of the sGC HD core highlights this discrepancy (Fig. 3F). In contrast, activation of sGC is accompanied with a large-scale conformational change, “straightening” the HD (Fig. 3H) and adopting the HD conformation that closely matches that of Cya (Fig. 3H). The position of the “kink” in the HD of sGC approximately corresponds to the membrane-cytosol interface in Cya. Thus, the very distant yet related proteins sGC and Cya (as well as AC9 and other membrane ACs) may be subject to very similar modes of regulation involving changes in the HD, which may result in changes in the catalytic domain of the protein. While in sGC the process is guided by the heme-containing receptor domain, in the membrane ACs this function is likely performed by the TM domain.

### The TM domain of Cya as a putative receptor module

The structure of Cya revealed several prominent cavities in the TM domain of the protein, which may serve a stabilizing or regulatory role (Fig. 4). A negatively charged cleft (site Ex1) is formed at the extracellular interface of the two 6-TM bundles (Fig. 4A, D, E). The negative charge of this pocket is provided by the residues D123, E164 and D170 of each monomer, facing into the cavity (Fig. 4E). This region may be involved in binding of positively charged ions, small molecules, lipids or peptides. The ability of NB4 nanobody to interact with this pocket spuriously indicates that it may also be a site of interaction with a yet unknown natural protein partner.

**Fig. 4.**
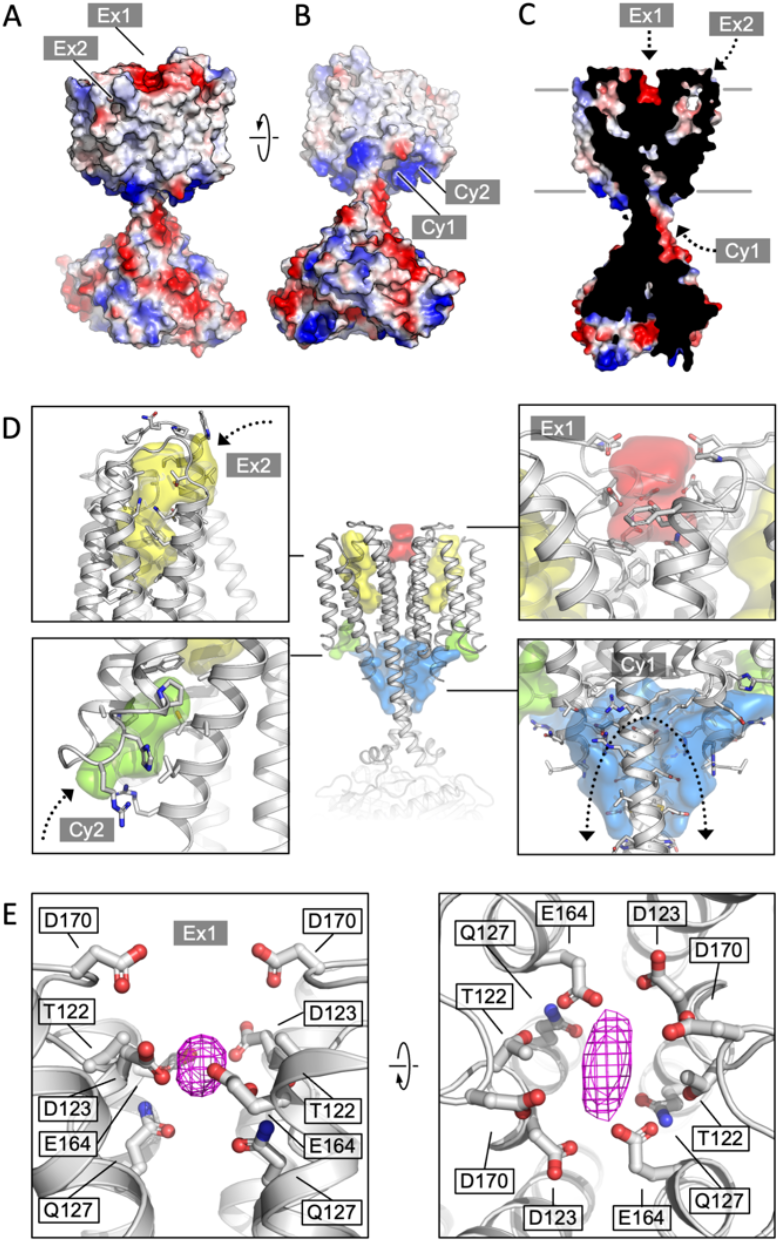
Cya TM domain as a receptor module. (**A**) A view of the Cya structure (nanobody NB4 not shown) in surface representation, coloured according to electrostatic potential. The location of two putative binding sites, sites Ex1 (negatively charge) and Ex2 (hydrophobic) are indicated. (**B**) Similar as (A), a view at the Cya structure from the cytosol, showing the locations of the positively charged sites Cy1 and Cy2. (**C**) A slice through the structure shows the internal cavities with access points Ex1, Ex2 and Cy1. (**D**) The density maps corresponding to the internal cavities within the TM region of Cya, calculated using 3V (*56*) and low pass-filtered to 3 Å for presentation purposes. Arrows indicate the access to the cavities. (**E**) A prominent density featured in the density map of Cya-NB4 complex, occupying the site Ex1. Polar and negatively charged residues surround the density, consistent with a binding site for metals (or organic cations).

Additionally, a prominent pocket open to the extracellular side of the protein is formed within each TM bundle (site Ex2; Fig. 4A, C-D). This pocket may accommodate small molecules or lipids, with a possible access route from the outer leaflet of the lipid bilayer surrounding Cya. A similar internal pocket is present in the TM1-6 bundle in AC9 (Fig. S8). Deep pockets on the cytosolic side of the Cya TM region are formed between the HD domain and the N-terminus/TM1 (site Cy1), as well as between TM1-3 (site Cy2; Fig. 4B-D). The entrances into these pockets are lined by positively charged residues R43, R44, R46 of the N-terminus, as well as R203 and R207 in the HD domain from the adjacent monomer (Fig. 4B-D). The positive charge of this region indicates a potential role in interactions with the phospholipid headgroups or positively charged peptides or small molecules. The interpretation of these cytosolic intramembrane pockets requires caution, as the residues 1-40 of Cya are not resolved in our 3D reconstruction but may interact with and occlude these pockets. Analysis of the sites Ex2-Cy2 shows that they are discontinuous, precluding formation of a channel traversing the entire width of the membrane (Fig. 4D). Our MD simulations confirmed the ability of water molecules to enter into the Ex2 and Cy2 site, but no transmembrane water transport could be observed (Fig. S9F). Thus, while the pockets Ex2 and Cy2 provide support to the hypothesis of the AC TM domain as a receptor, any translocation events would have to involve substantial conformational rearrangements opening the connection between the two sites.

Our density map features a small but prominent density in the site Ex1 (Fig. 4E, Fig. S6), which presently can not be assigned to a specific entity, but which could correspond to a bound metal (Na^+^, Mg^2^, Mn^2^ or a yet unknown component co-purified with the protein from *E. coli*). It is conceivable that disruption of this negatively charged interface may lead to the loss of the rail structure of the TM region, with concomitant changes in the HD helix arrangement and ultimately the catalytic domain of Cya. To test the behavior of this site, we performed molecular dynamics (MD) simulations (Fig. S9). The site Ex1 behaved as a genuine metal binding site, occupied by the K^+^ and Mg^2+^ions similarly to the metal binding site in the Cya catalytic center over the course of the simulation (Fig. S9B-E). Thus, the evidence obtained experimentally and using MD simulations strongly supports a function of the extracellular surface of Cya as a receptor for positively charged ligands.

### The Ex1 site controls the helical and catalytic domain

To test the role of the Ex1 site experimentally, we mutated the polar residues lining this site (T122, D123, Q127, E164 and D170) to Ala (Fig. 5A-C). The resulting construct (referred to as Ex1-5A, Fig. 5A, C) was successfully expressed and purified in digitonin (Fig. S10A). The thermostability profiles of the Ex1-5A mutant in the absence or in the presence of a nucleotide were similar to those of the wild-type Cya (Fig 5D, Fig. S10B-C). In contrast, the enzymatic activity of Ex1-5A was substantially reduced (with a dramatic increase in apparent K_m_, Fig. 5E), with a 4-fold reduction in apparent affinity for a nucleotide inhibitor, MANT-GTP (judged by the MANT-GTP IC_50_ values, Fig. 5F).

**Fig. 5.**
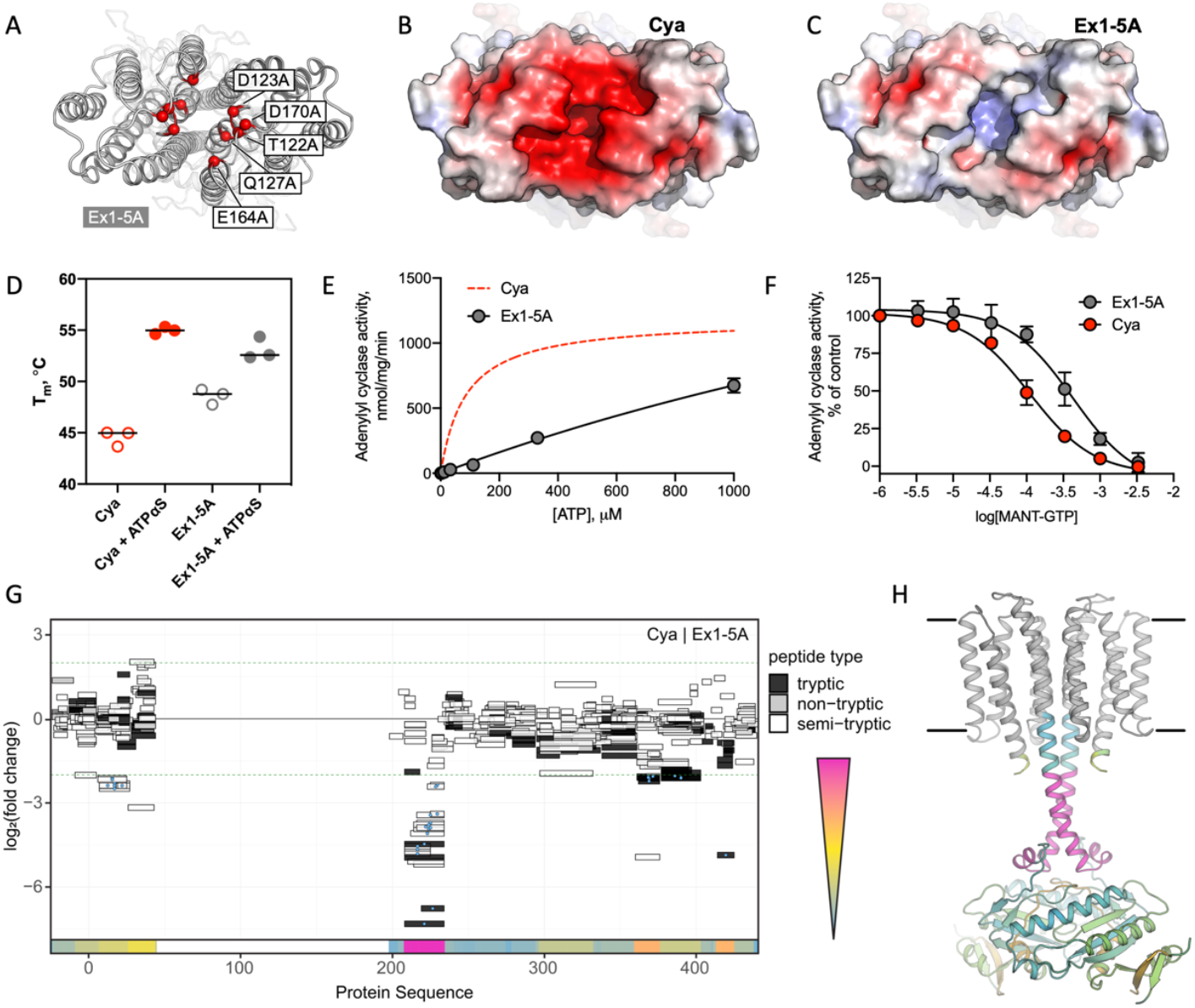
Extracellular site Ex1 is linked to the adenylyl cyclase activity of Cya. (**A**) An illustration of the Ex1 site residues mutated to generate the Ex1-5A mutant, substituting the five indicated residues with Ala. (**B**) Calculated electrostatic potential of the wild-type Cya. (**C**) Same as B, for the Ex1-5A mutant. (**D**) The mutant shows thermostability profile consistent with that of the wild-type protein, based on the observed T_m_ values in the presence and in the absence of a nucleotide analogue. For experiments in D-F, n = 3; data are shown as mean ± S.E.M. (**E**) The enzymatic properties of the mutant are substantially affected by the mutation (the dashed red curve corresponds to the fit shown for Cya in Fig. 1D for comparison). (**F**) The affinity of the Ex1-5A mutant for MANT-GTP is reduced (110 and 420 μM, respectively). (**G**) Limited proteolysis-coupled mass spectrometry (LiP-MS) analysis of Cya and Ex1-5A mutant. The graph indicates sequence coverage and the identified tryptic, semi-tryptic or non-tryptic peptides. Significantly changing peptides (|log2(FC)| > 2; q-value < 0.001) are marked with a blue dot. A bar within the plot is coloured according to the change in protease accessibility at each peptide (blue = no change, pink = high fold change; absolute log2 transformed fold changes range from 0 to 7.3). (**H**) A model of Cya coloured according to the bar in (E).

To investigate the effects of the Ex1-5A mutant on Cya structure, we performed limited proteolysis–coupled mass spectrometry (LiP-MS) experiments on both the wild-type and the mutant in the absence of added nucleotides, and compared the peptides obtained by pulse proteolysis with proteinase K (PK) (Fig. 5G). Comparative analysis of the LiP-MS profile of the wild-type Cya and the Ex1-5A mutant revealed a significant increase in protease accessibility of the HD in the mutant (Fig. 5G-H). Thus, modification of the extracellular site Ex1 of Cya leads to changes in the dynamics of its cytosolic HD, accompanied by a dramatic reduction in enzymatic activity.

## Discussion

The cryo-EM structure of *M. tuberculosis* Cya provides a unique insight into the assembly and regulation of a model membrane AC. To this day, the functional role of the TM region in the polytopic membrane AC, such as Cya or the mammalian AC1-9, remains elusive. Why does a cell need an AC with such an elaborate membrane anchor? A lipid anchor or a single TM helix would be sufficient to target the catalytic domain to the membrane compartment where cAMP production is required. Our structure provides two possible reasons for the ACs to have such a TM domain: (i) to facilitate the assembly of the HD domain, (ii) to act as a receptor module, binding ligands at several newly identified putative ligand binding sites, including the pockets within the Cya monomers and at the Cya-Cya interface (Fig. 6). The two functions are not mutually exclusive, as the ligand interactions with the membrane region of the AC may influence the HD assembly and thus regulate the cyclase function. Previous work utilizing chimeric constructs composed of fragments of the quorum sensing receptor CqsS from *Vibrio harveyi* and Cya (*20*) or mammalian membrane ACs (*28*) revealed that the membrane anchors of the ACs may act as orphan receptors for yet unknown ligands. Together with the proposed functional coupling between the TM domain and the catalytic site of Cya, the structure described here is consistent with these findings, offering molecular insights into the potential receptor role of the membrane anchor of a model membrane AC.

**Fig. 6.**
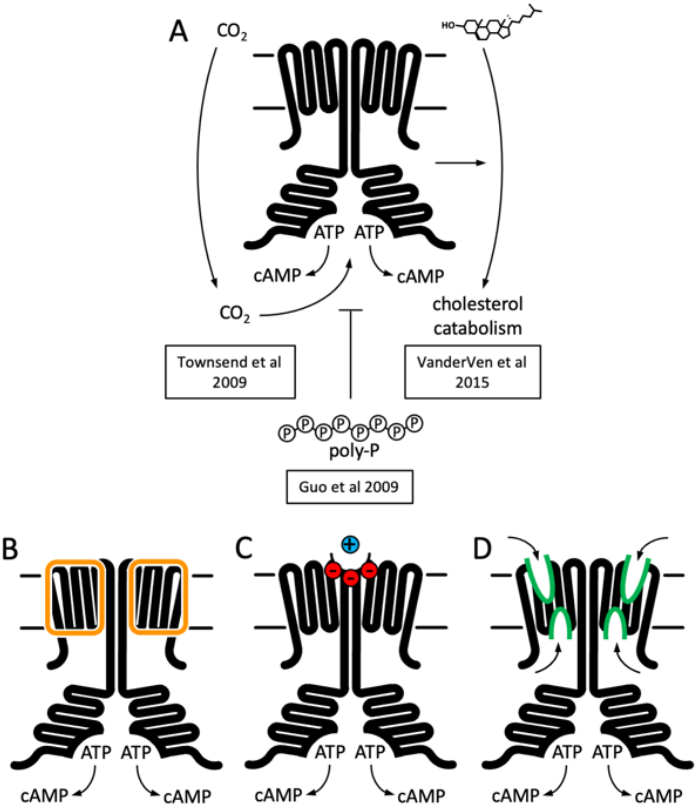
Function and structure of Rv1625c / Cya. (**A**) Known regulators and cellular functions of Cya. (**B-D**) Insights into the function of the membrane domain of Cya, with new functions of the TM domain suggested by the cryo-EM structure: a stabilizer of the cytosolic domain assembly (B), a receptor for positively charged agents via the Ex1 site at the Cya dimer interface (C), a receptor of yet unknown ligands via sites Ex2 / Cy1 / Cy2 (D).

Our experimental results and simulations point to a possible link between the enzymatic activity of Cya and binding of small cations (such as metals ions) to its Ex1 site. It is possible that transient interactions of cations with protein surfaces play a basic role in surface charge compensation. However, the properties of the Ex1 site are strongly suggestive of a specific ligand binding site, with a cluster of ten polar residues (six negatively charged residues) at the Cya dimer interface pointing towards the Ex1 cavity. The most telling evidence for the role of this site in cation binding is the MD simulation, which revealed two locations on the Cya surface where the metal ions dwell: the established metal binding site in the catalytic pocket, and the site Ex1. Modification of the charges of the Ex1 by mutagenesis reduced the activity of the protein, further suggesting that this site at the extracellular surface of the protein likely plays a pivotal role in controlling the assembly of the catalytically active dimeric Cya.

The presence of potential ligand binding pockets at the extracellular surface of Cya lends strong support to the long-standing idea that the TM domains of the ACs may act as receptor modules for yet unknown ligands (*20*). The possibility of direct regulation of cAMP production via the membrane anchors of the ACs would have long-reaching consequences: a vast repertoire of pharmacological agents on the market today act via GPCRs coupled to membrane ACs (*29*). Direct modulation of the cAMP production through the AC membrane domains could revolutionize the approaches to drug development for a wide range of diseases where the GPCRs are currently the primary drug targets. A related notion of importance for molecular pharmacology and medicinal chemistry is the potential interactions between the already existing drugs and the membrane domains of the ACs. Such interactions may lead to unwanted side-effects associated with cAMP signaling, such as emesis and changes in heart rate and contractility (*30*). An example of this is the antifungal drug miconazole, which is known to have cardiotoxic effects (*31, 32*). Miconazole has recently been shown to directly activate AC9, likely via its TM domain (*33*). The interaction with AC9 may contribute to the cardiorespiratory side-effects of this drug. Similar interactions involving other drug / AC combinations may have to be systematically evaluated, especially in the cases where changes in cAMP levels or in the downstream signaling events are recognized as side-effects.

In the absence of known interaction partners for the membrane regions of the ACs it is difficult to predict the effect of ligand binding to any of the pockets we find in the Cya structure. Nevertheless, the structure hints at ways that could be exploited by various agents to affect the AC activity via the membrane domain. The closest example of cyclase modulation through the receptor-mediated effect on the HD helices is the case of soluble GC, described at atomic resolution, has been detailed above (Fig. 3) (*23*). The presence of disease-linked mutations in the HD regions of AC5 (*34, 35*) and retGC1 (*36, 37*) underscore the importance of this domain for cyclase structure and function. It is clear that the HD region plays a vital part in AC and GC assembly and stability. This is evident from our experiments with the *M. intracellulare Cya* (*17*), as well as the results of others using Cya and mammalian ACs as model enzymes (*10, 26, 28*). It remains to be determined whether any agents can elicit conformational changes in the membrane domain of Cya (or in any of the mammalian membrane ACs), leading to substantial changes in the HD similar in scale to the changes observed in the sGC during its activation. Our MD simulations and the experiments with the Ex1-5A mutant of Cya are suggestive of a receptor-transducer-catalyst relay, where the extracellular portion of the TM region acts as a “receptor” for a yet unknown ligand, the HD transduces the activation, and the catalytic domain catalyses ATP to cAMP conversion. This notion is further supported by previous work on Cya that identified the HD region as a transducer of a putative signal (*20, 26*). The structure of Cya can serve as a starting point for exploration of the TM domain-mediated regulation of membrane ACs.

## Materials and Methods

### Protein expression and purification

#### Expression and purification of the full-length Cya

Cya cloned into a vector with an N-terminal strep tag and a 3C cleavage tag was expressed in *E. coli* BL21(DE3)RIPL cells grown in TB medium. Protein expression was induced when at OD600 of 3.0 using 0.3 mM IPTG. After 3 hours of induction the cells were harvested. The membranes were prepared using cells lysis by three passes in Emusiflex high pressure homogenizer in a buffer containing 50 mM Tris pH 7.5, 200 mM NaCl, 5 μg/ml DNase and 1 mM PMSF. Lysed cells were centrifuged at 12000 rpm using a Ti45 rotor for 30 mins. The resulting supernatant was spun down by ultracentrifugation using Ti45 rotor at 40000 rpm for 1 hour, re-suspended in a buffer containing 50 mM Tris pH 7.5 and 200 mM NaCl and ultracentrifuged again. The resulting membrane pellet was resuspended in the same buffer, flash-frozen and stored at −80°C until purification.

For purification, the membranes were thawed and resuspended in a buffer containing 50 mM Tris pH 7.5, 200 mM NaCl, 10% glycerol and 1% sol-grade dodecylmaltoside (DDM, Anatrace), mixed at 4°C for 1 hour and ultracentrifuged. The supernatant was incubated with Strep-tactin superflow resin for 1 hour at 4°C. The resin was washed with a volume 25 times that of the resin bed of a buffer containing 0.1% digitonin and the eluted with 5 mM desthiobiotin. The eluted protein was concentrated and injected onto Superose 6 Increase column pre-equilibrated with a buffer containing 50 mM Tris pH 7.5, 200 mM NaCl 0.1% digitonin and 10% glycerol. For Cryo-EM samples glycerol was omitted during size exclusion chromatography step.

#### Cya-SOL expression and purification

Cya-SOL construct was generated by cloning the sequence encoding the Cya residues 203-433 into a vector with an N-terminal 10xHis tag followed by a 3C cleavage site. The construct was expressed in *E. coli* BL21(DE3)RIPL cells grown in TB medium. Expression was carried out under conditions similar to those used for expression of the full-length protein, with a 5-hour induction at 20°C. The cells were collected by centrifugation, lysed and the cleared lysate was incubated with Ni-NTA resin for 1 hour. The resin was washed with a volume 15 times the resin bed volumne of wash buffer containing 50 mM Tris pH 7.5, 200 mM NaCl, 10% glycerol and 20 mM imidazole, followed by an additional wash step with a volume of 25 times the volume of the resin bed volume with a buffer containing 50 mM imidazole. The protein was eluted with a buffer containing 250 mM imidazole, concentrated and desalted using a GE PD-10 Sephadex G-25 desalting column. The protein was mixed with 3C protease (1/50 w/w) and incubated at 4°C overnight. The protein was passed through pre-equilibrated Ni-NTA resin to remove the 3C protease and purified by SEC using Superdex 200 Increase column.

### Nanobody library generation and selections

To generate desired immune response in heavy chain-only IgG subclass, an alpaca was immunized four times in two-week intervals, each time with 200 µg purified Rv1625c in PBS containing 0.02% (w/v) β-DDM. The antigen was mixed in a 1:1 (v/v) ratio with GERBU Fama adjuvant (GERBU Biotechnik GmbH, Heidelberg, Germany) and injected subcutaneously in 100 μL aliquots into the shoulder and neck region. Immunizations of alpacas were approved by the Cantonal Veterinary Office in Zurich, Switzerland (animal experiment licence nr. 172/2014). One week after the last injection, 60 mL of blood was collected from jugular vein for isolation of lymphocytes (Ficoll-Paque® PLUS, GE Healthcare Life Sciences, and Leucosep tubes, Greiner). Approx. 50 mio. cells were used to isolate mRNA (RNeasy Mini Kit, Qiagen) that was reverse transcribed into cDNA (AffinityScript, Agilent, US) using the gene specific primer. The VhH (nanobody) repertoire was amplified by PCR and phage library was generated by fragment exchange cloning (*22*) into a PmlI-linearized pDX phagemid vector. The resulting VhH-phage library (size 4.5 e6) was screened by biopanning against the immobilized target. For that purpose VI23.60 containing Strep-tag® was immobilized on the Strep-Tactin® coated microplate (IBA Lifesciences GmbH, Germany) and three rounds of selection were performed. 195 single clones from the enriched nanobody library were induced to express polyhistidine-tagged soluble nanobodies in the bacterial periplasm and analysed by ELISA for binding to the target. 96 ELISA-positive clones were Sanger sequenced and grouped in 17 families according to their CDR3 length and sequence (*22*).

### Nanobody expression and purification

Nanobody NB4 was expressed in BL21(DE3)RIPL cells in TB medium supplemented with 2 mM magnesium chloride and 0.1% glucose by induction at an OD600 of 0.7 using 1 mM IPTG at 26°C for 16 hours. The periplasmic fraction was isolated by resuspending the cell pellet in 2.5x w/v cold TES buffer (200 mM Tris pH 8.0, 0.5 mM EDTA and 0.5 mM sucrose and 1 mM PMSF) for 45 mins, followed by an overnight incubation with twice the amount of a 4-fold diluted TES buffer. The suspension was spun down and the supernatant was used for protein purification with Ni-NTA resin, following the same procedure as that used for Cya-SOL. The eluted nanobody was concentrated and further purified using SEC with a Superdex 200 Increase column.

### Adenylyl cyclase activity assay

Adenylyl cyclase activity assays were performed as described previously (*17*). In brief, the assay was carried out in a reaction volume of 200 μl with 50mM Tris pH 8.0, 200 mM NaCl, 5 mM MgCl2 5 mM MnCl2 and 0.1% digitonin. For determination of Km, ATP concentration was varied from 0 to 1000 mM in presence of 10 nM [^3^H]ATP (PerkinElmer). The reaction was initiated by adding ATP to the reaction solution containing 0.005 mg/ml Cya, 0.0075 mg/ml NB4, followed by an incubation for 10 min at 30°C. The reaction was stopped by incubating the reaction mixture at 95°C for 4 minutes and by addition of 20 μl of 2.2 M HCl. The stopped reactions were applied to 1.3 g of aluminum oxide in disposable columns. The cAMP was eluted with 4 ml of 100 mM ammonium acetate into scintillation vials and mixed with 12 ml scintillation liquid (Ultima Gold). The amount of radioactive cAMP was measured using a liquid scintillation counter. The activity assays for Cya-SOL were performed identically, using 0.08 mg/ml Cya-SOL in an assay.

### Isothermal titration calorimetry

The isothermal titration calorimetry (ITC) experiments were performed using a Microcal ITC200 instrument with cell temperature maintained at 25°C and with stirring set to 750 RPM. In total, 15 injections were performed per experiment with each injection set at 2 μl and a pre-injection volume of 0.8 μl. Cya-SOL was kept in the cell at a concentration of 30 μM and NB4 was kept in the syringe at 300 μM. All ITC measurements were performed in triplicates. The results were analyzed using sedphat and NITPIC. The figures describing the ITC results were generated using GUSSI (*38*).

### Protein thermal unfolding

Protein thermal stability was measured using nanoDSF on a Prometheus panta instrument (NanoTemper) (*39*). The protein was measured at 0.5 mg/ml concentration in a buffer containing 50mM Tris pH 7.5, 200 mM NaCl and 0.1% digitonin, using NT.48 capillaries. For samples with ligand, 1mM ATPαS and 5 mM MgCl2 was added and allowed to incubate at RT for 10 minutes. The samples were spun at 13000 g on a tabletop centrifuge for 1 minute before measuring. Thermal unfolding experiments were carried out at a temperature increment of 1°C/min in triplicates. Tm was calculated as the first derivative of intrinsic protein emission ratio at 350 nM and 330nM using PR.Panta analysis software.

### Cryo-EM sample preparation

For Cryo-EM sample preparation, freshly purified full-length Cya in 0.1% digitonin was concentrated and mixed with NB4 at a molar ratio of 1:1.5. Additionally, 5 mM MnCl2 and 0.5 mM MANT-GTP were added and the mixtures were incubated on ice for 30 minutes. The final concentration of Cya was 5-6 mg/ml. An aliquot of 3.5 μl of sample was placed on the glow-discharged cryo-EM grid (Quantifoil R1.2/1.3 or Quantifoil R2/1), blotted and plunge-frozen in liquid ethane using a Mark VI Vitrobot instrument maintained at 100% humidity with blot force 20 and blot time of 3 seconds. The grids were cryo-transferred for storage in liquid nitrogen.

### Cryo-EM data acquisition and image analysis

The cryo-EM data were obtained at the SCOPEM facility at ETHZ using a 300 kV Titan Krios electron microscope (FEI) equipped with a K3 direct electron detector with a pixel size of 0.33 Å/pix (in super-resolution mode), at a defocus range of −0.5 to −3.0 μm. All movies were dose fractionated into 40 frames. The movies for dataset 1 were recorded with a total dose of 54 e-/Å^2^, dataset 2 -with a dose of 47 e-/Å^2^, and for dataset 3 - a dose of 44 e-/Å^2^.

All data processing was performed in relion 3.0 (*40*). All micrographs were motion corrected using motioncorr 1.2.0 (*41*) and binned two-fold. All micrographs were CTF corrected using Gctf (*42*). Particles were autopicked using templates from manual picking. In total 1692104 particles were picked for data set 1, 1898968 particles for data set 2 and 990286 particles for data set 3. After several rounds of 2D classification, data set 1, 2 and 3 were left with 253789, 1173076 and 741081 particles respectively. 3D Classification with four classes was used to further process each dataset, with C1 and C2 symmetry imposed. The particles from the best classes in each data set were chosen and further refined. The extracellular density for NB4 was masked out for all subsequent refinements to generate refined 3D maps at a resolution of 4.37 Å (dataset 1), 4.25 Å (dataset 2) and 4.61 Å (dataset 3) in C2 symmetry. The particles were merged into a single selection and subjected to refinement, ctf refinement and particle polishing, yielding a final refined map of 3.57 Å resolution (C2 symmetry). The same particle selection produced a reconstruction at 3.83 Å resolution without symmetry imposed (C1). Model building was performed in coot (*43*). The model was refined using phenix.real_space_refine in Phenix (*44*). For model validation the model atoms were randomly displaced (0.5 Å), and the resulting model refined using one of the refined half maps (half-map1). Map vs model FSC was calculated using the model against the corresponding half-map1 used for refinement, and for the same model versus the half-map2 (not used in refinement) (*45*). Model geometry was assessed using MolProbity (*46*).

### Protein crystallization, X-ray data collection, processing and structure determination

Crystallization of Cya-SOL-NB4 complex was performed using standard vapour diffusion techniques at 20°C. Concentrated protein complex was prepared by mixing purified Rv1625c and NB4 in 1:1.2 ratio in a buffer of the following composition: 20 mM Tris-HCl pH 7.5, 150 mM NaCl, 5 mM MnCl2 and 1 mM MANT-GTP. This protein solution was used to set up 96-well sitting drop crystallization trials using TPP Mosquito LCP robot. Formulatrix Rockimager was used to visualize crystal formation. Multiple crystal hits were obtained, and selected conditions were used as starting points for further crystal optimizations. The optimal crystals were obtained after mixing 1.5 μl of protein (20 mg/ml) with 1.5 μl of reservoir solution (0.1 M Na-acetate pH 5.5, 0.02 M CaCl2, 30% MPD). The crystals were gently transferred to cryoprotectant consisting of 0.05 M Na-acetate pH 5.5, 0.01 M CaCl2, 35% MPD, and crystals were subsequently mounted onto crystal loop (Hampton Research) and flash frozen in liquid nitrogen.

X-ray data collection was performed at the PXI and PXIII beamlines at the Swiss Light Source synchrotron in Villigen, Switzerland. The best dataset was collected to 1.97 Å resolution from a single crystal at cryogenic temperature (100 K) using Eiger detector with oscillation range of 0.1°. The data was processed using MOSFLM and XDS (*47, 48*). The resolution cutoff was chosen taking into account the values of CC1/2 and mean I/sigma(I) (*49*). Phasing/refinement was performed using Phenix (*44*). Phases were resolved by molecular replacement (MR) using templates (cya: 5O5K, nanobody: 6FPV). Coot was used for model building and geometrical optimization (*43*). Crystallographic data collection and refinement statistics are shown in Table S2.

### Molecular dynamics (MD) simulations

The MD simulations were performed using CHARMM36m force field in GROMACS 2019.3 (*50*). The missing N-and C-terminus of Cya were modelled using I-TASSER server (*51*), and the protein was inserted into lipid membrane, solvated and ionized using the Membrane Builder tools in CHARMM-GUI (*52*). Lipid membrane composed of 60 POPC, 40 POPG, 40 POPE, 20 POPI, 20 DLGL and 20 cholesterol molecules. The system was solvated with TIP3 water molecules extended to 25 Å from the edge of the protein and with per lipid hydration number of 50. Subsequently the system was neutralized with Cl^-^ ions, and then was brought to a final concentration of 0.15 M KCl or 0.15 M MgCl2. All of the generated systems were subjected to energy minimization, and 6-step NVT and NPT equilibration using the default scheme provided in the CHARM-GUI (3), followed by 200 ns of a production run. Final trajectories were analyzed using tools in available the GROMACS package. The Volmap tool in VMD (*53*) was used for generating occupancy/density maps. The figures related to the MD simulations were prepared using VMD or Pymol.

### Limited proteolysis-coupled mass spectrometry

Wild-type and mutant Cya protein preparations were diluted in LiP buffer (1 mM MgCl2, 150 mM KCl, 100 mM HEPES-KOH pH 7.4) with 0.1% digitonin. Mutant and wild-type samples were split into 8 samples each at a protein amount of 2 µg of protein per 50 µL of buffer. Four out of eight WT and mutant samples were treated with proteinase K from *Tritirachium album* (Sigma Aldrich) (limited proteolysis, LiP), whereas the other four were treated with water instead (TC). The samples were incubated in a Thermocycler for 5 minutes at 25°C. Proteinase K was inactivated by heating the samples to 99°C for 5 minutes, then incubating them at 4°C for 5 minutes, followed by the addition of the same volume of 10% sodium deoxycholate. LiP and TC underwent the same procedures.

#### Tryptic digest

Following the addition of sodium deoxycholate, disulfide bonds were reduced by adding tris(2-carboxyethyl)phosphine to a final concentration of 5 mM and incubating the samples at 37°C for 40 minutes with slight agitation. Free cysteine residues were alkylated with iodoacetamide at a final concentration of 40 mM and 30 minutes of incubation at room temperature in the dark with slight agitation. The samples were diluted with 100 mM ammonium bicarbonate to a final sodium deoxycholate concentration of 1 %. Lysyl endopeptidase LysC was added at an enzyme to substrate ratio of 1:50 and samples were incubated for one hour at 37°C with slight agitation. Next, trypsin was added at an enzyme to substrate ration of 1:50 and incubated at 37°C overnight with slight agitation. The digestion was stopped by adding 50% formic acid to the samples to achieve a final concentration of 2% formic acid. Precipitated sodium deoxycholate was removed by filtering through a Corning® 2 µM PVDF plate and samples were further desalted on a 96-well MacroSpin plate (The Nest Group). Peptides were eluted with 80% acetonitrile, 1% formic acid and dried in a vacuum centrifuge. After drying, samples were reconstituted in 20 µL 0.1% formic acid and iRT peptides (Biognosys) were added.

#### LC-MS/MS data acquisition

Samples were measured on an Orbitrap Exploris™ 480 mass spectrometer (Thermo Fisher), equipped with a nanoelectrospray source and an Easy-nLC 1200 nano-flow LC system Thermo Fisher). 1 µL of digest was injected and separated on a 40 cm x 0.75 i.d. column packed in-house with 1.9 µm C18 beads (Dr. Maisch Reprosil-Pur 120) using a linear gradient from 3 to 35 % B (eluent A: 0.1% formic acid, eluent B: 95% acetonitrile, 1% formic acid). Gradient duration was 30 minutes, whereas the whole method was 60 minutes long. Samples were measured at a constant flowrate of 300 nL/min while the column was heated to 50°C. All samples were acquired in DIA (41 windows, 1 m/z overlap) and analyzed in Spectronaut v15 (Biognosys). Further data analysis was carried out in R using mainly the R package protti (*54*). Briefly, the abundances of Rv1625c mutant and wild type where compared in the tryptic controls and the protein abundances in the LiP-samples were corrected accordingly. Statistical testing on peptide level to detect peptide abundance differences was conducted employing the proDA (*55*) algorithm, implemented in protti. The Rv1625c PDB file was edited and the b-factors were replaced with the maximum absolute value of the calculated log2(fold change) at each position. In pyMOL the protein was then colored according to the replaced b-factors to highlight regions changing regions.

#### LiP-MS data interpretation

Whether a peptide decreases or increases in abundance is dependent on the accessibility of the native protein to PK. In a standard LiP-MS experiment, three peptide types can be detected, dependent on the proteolytic cleavage:

- Semi-tryptic peptides are generated by a cleavage of PK on either the N-terminal or the C-terminal side of the peptide and a cleavage by trypsin on the respective other side.
- Tryptic peptides are not cleaved by PK at all.
- Non-tryptic peptides were cleaved by PK on both sides.

Depending on the peptide type an increase or decrease in abundance can be interpreted in different ways. A tryptic peptide that decreases in abundance was additionally cleaved by PK, hence it disappears. This likely means that the protein region became more accessible to PK. On the other hand, a tryptic peptide that decreases in abundance can be interpreted as the region becoming less accessible to PK. A semi-tryptic peptide that increases in abundance can be explained as the protein region cleaved by PK becoming more accessible. A semi-tryptic peptide that decreases in abundance can be explained in two different ways: either the protein region became more protected, hence inaccessible to PK, or the protein region became more accessible and the peptide was not detected because of additional PK cleavage sites that were introduced with the conformational change.

## Acknowledgments

We thank Emiliya Poghosyan and Elisabeth Müller-Gubler (EM Facility, PSI), and Miroslav Peterek (ScopeM, ETH Zurich) for their support in cryo-EM data collection. We also thank Spencer Bliven and Marc Caubet Serrabou (PSI) for support in high performance computing.

## Funding

Swiss National Science Foundation (150665; VMK)

Swiss National Science Foundation (176992; VMK)

Swiss National Science Foundation (184951; VMK)

Vontobel Stiftung (VMK)

## Author contributions

Conceptualization: VM, VMK

Methodology: VM, BK, DS, IV, SS, PP, VMK

Investigation: VM, BK, AK, DS, IK, TI, IV, VMK

Visualization: VM, BK, DS, VMK

Funding acquisition: VMK

Project administration: VMK

Supervision: PP, VMK

Writing – original draft: VM, VMK

Writing – review & editing: VM, BK, DS, PP, VMK

## Data and materials availability

The atomic coordinates and structure factors have been deposited in the Protein Data Bank (7YZ9, 7YZI, 7YZK); the density maps have been deposited in the Electron Microscopy Data Bank (EMD-14388, EMD-14389). The mass spectrometry data will be deposited at ProteomeXChange via PRIDE. All other data are available in the main text or the supplementary materials.

## Supplementary Materials

Figs. S1 to S10

Tables S1 to S2

