## Supplementary Materials for "Structure of *Mycobacterium tuberculosis* Cya, an evolutionary ancestor of the mammalian membrane adenylyl cyclases"

**Supplementary Figures**

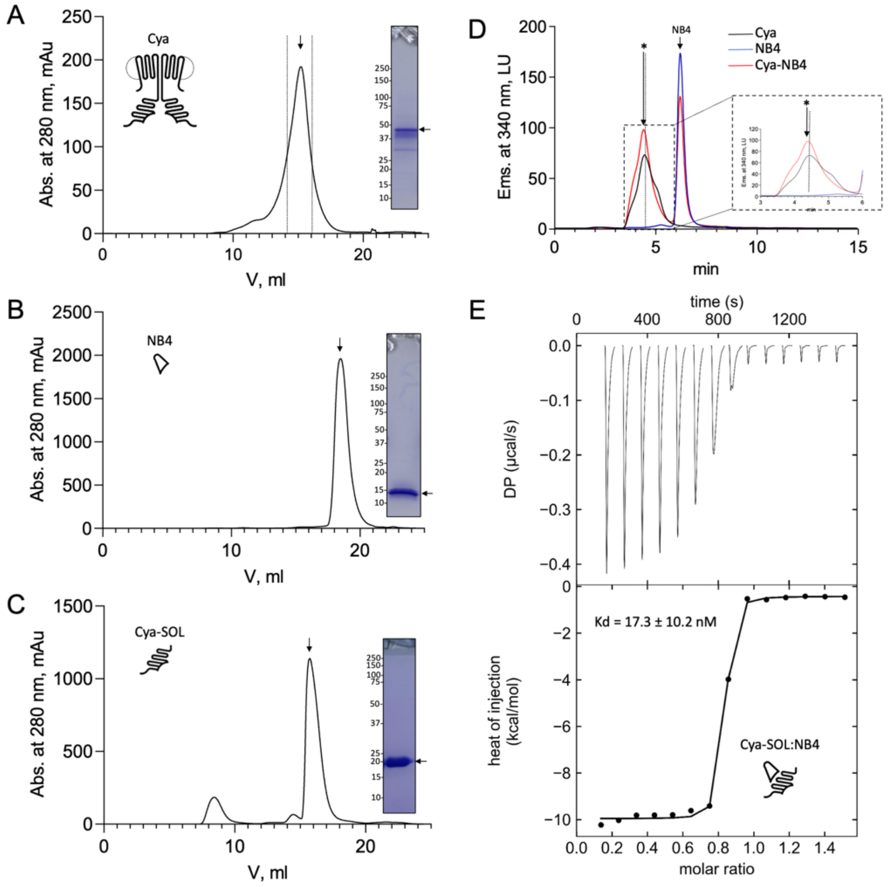


**Figure S1. Purification and characterization of Cya. (A)** Size-exclusion chromatography (SEC) and SDS PAGE of the purified Cya. SEC was peformed using a Superose 6 Increase column. **(B-C)** Same as in A, for NB4 (B) and for Cya-SOL. SEC was performed using a Superdex 200 Increase column. **(D)** Analytical SEC analysis Cya-NB4 interaction was performed using the Agilent Bio SEC-5 column. The shift of the SEC peak upon NB4 binding is indicated with an arrow; a dashed line indicates the position of the Cya alone. **(E)** Isothermal titration calorimetry (ITC) analysis of Cya-SOL / NB4 binding; mean $K_d$ ± S.D. is indicated in the graph (n = 3). ITC thermogram integration was carried out using NITPIC. Global analysis of integrated thermograms was performed using SEDPHAT and figures were generated using GUSSI.

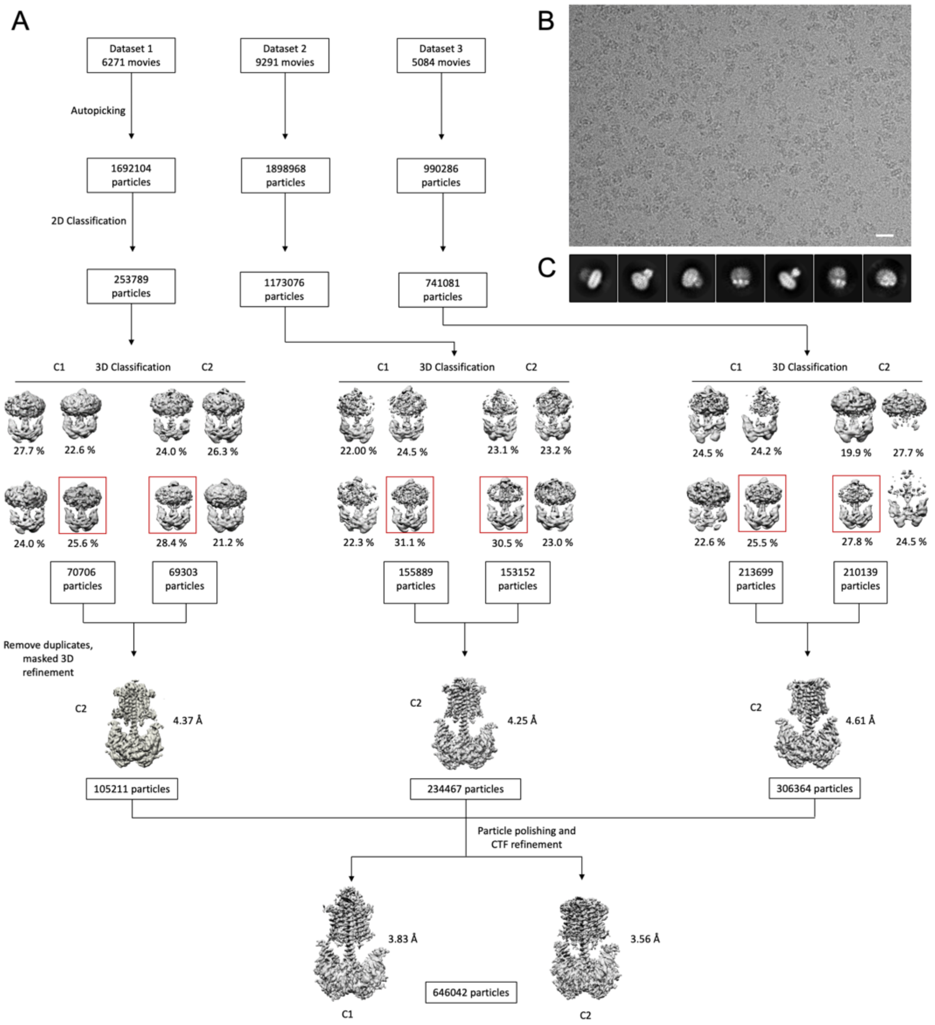


**Figure S2. Cryo-EM processing workflow. (A)** A processing pipeline for 3D reconstruction of the Cya-NB4 complex. Three data sets were processed in both C1 and C2 symmtery in parallel. Particles resulting from the best 3D classes were pooled, duplicates removed, and the resulting particle selection was refined in C2 symmetry for each data set. All three datasets were combined and further processed by particle polishing and CTF refinement, resulting in two density maps: one processed in C1 symmetry (at a resolution of 3.83 Å) and another one in C2 symmtery (at a resolution of 3.57 Å), as detailed in "Materials and Methods". **(B)** An example of a cryo-EM micrograph the bar corresponds to 200 Å. **(C)** Representative 2D classes reveal distinct views of the Cya-NB4 complex.

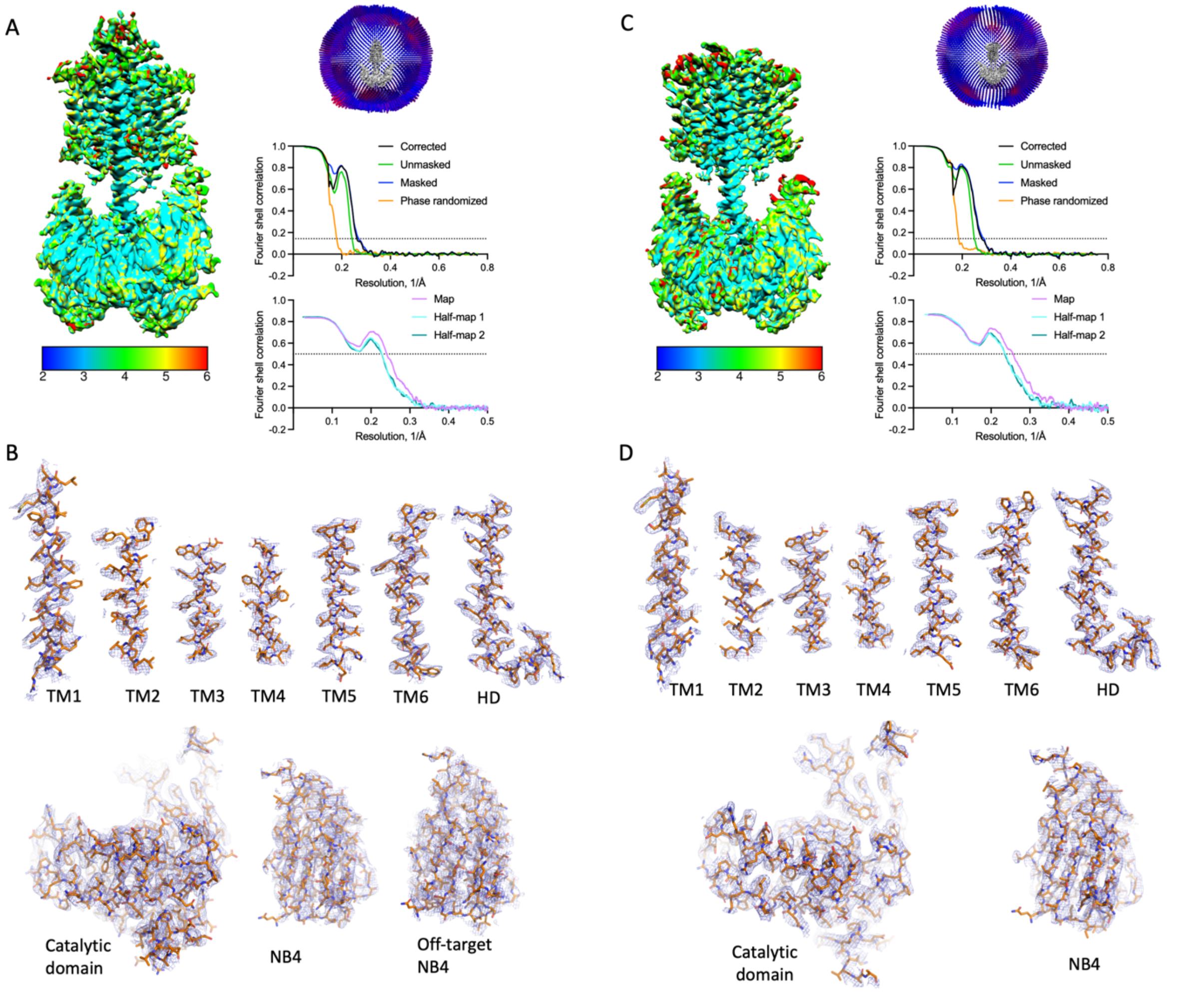


**Figure S3. Properties of the Cya-NB4 density maps. (A)** Local resolution, angular distribution and FSC curve of the Cya-NB4 complex processed in C1 symmetry. Scale bar indicates local resolution range in Å. **(B)** Isolated cryo-EM density for each of TM helix and the HD contoured at 8σ (*top*). Isolated Cryo-EM density for catalytic domain and NB4 contoured at 8σ, with extracellular NB4 contoured with 4σ (*bottom*). **(C-D)** Same as A-B, for the Cya-NB4 processed in C2 symmetry. All density map features are contoured at 8σ.

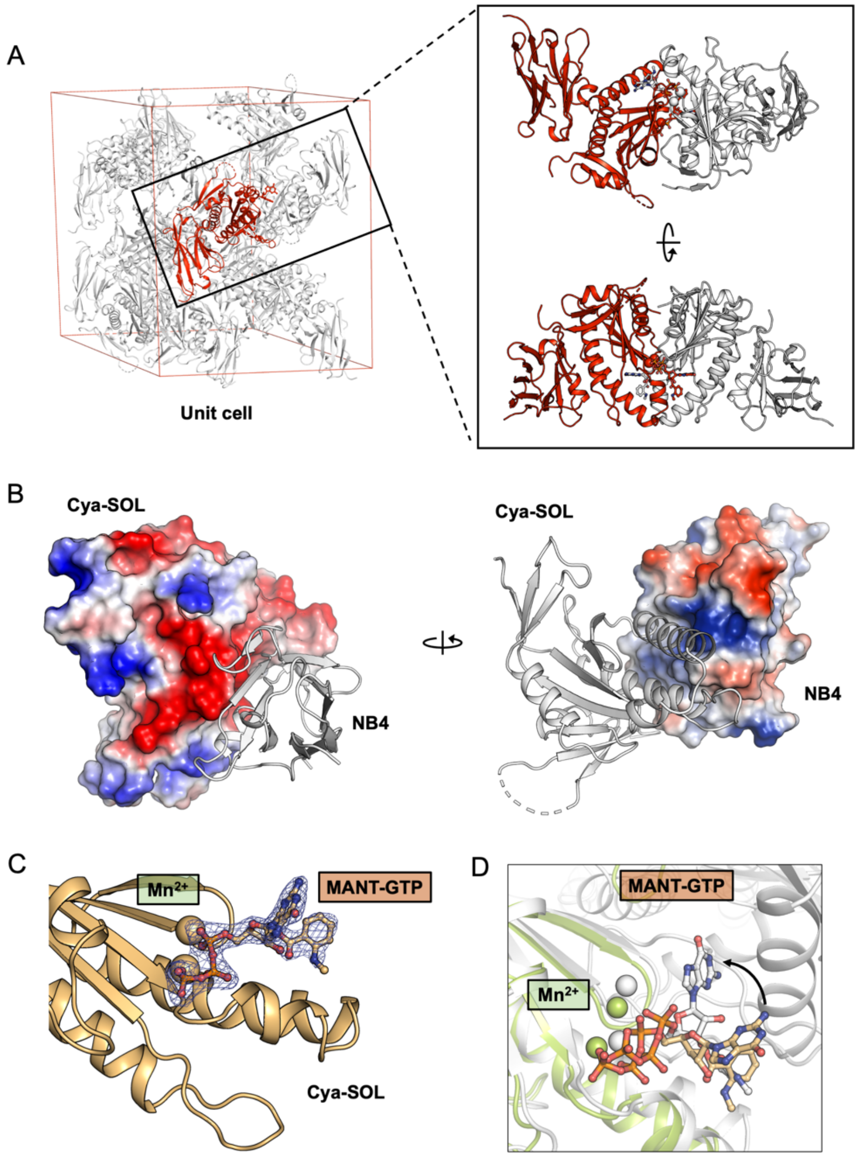


**Figure S4. X-ray structure of the catalytic domain of Cya, Cya-SOL, bound to NB4. (A)** Illustration of a crystallographic unit cell in the crystals of catalytic domain of Cya-NB4 complex showing the arrangement of one copy of Cya-SOL-NB4 complex (red) in an asymmetric unit (red). Analysis of the interaction between Cya-SOL-NB4 with its symmetry mates (grey) shows that Cya-SOL-NB4 does engages in non-native interactions with its neighbours, consistent with formation of a crystallographic dimer (*right*). **(B)** The surface representations of the catalytic domain of Cya (*right*) and NB4 (*left*), coloured according to the calculated electrostatic potential, suggest a role of the surface-exposed charged residues in the Cya-NB4 interaction. **(C)** Fo-Fc omit map of MANT-GTP (*left*) bound to Cya-SOL. Density is contoured at 3σ, carved to 1.6 Angstrom in Pymol. **(D)** Despite retaining the ability to bind the MANT-GTP molecule via one-half of its catalytic site, the position of the nucleotide is distinct from that observed in the structure of the nucleotide-bound dimeric Cya.

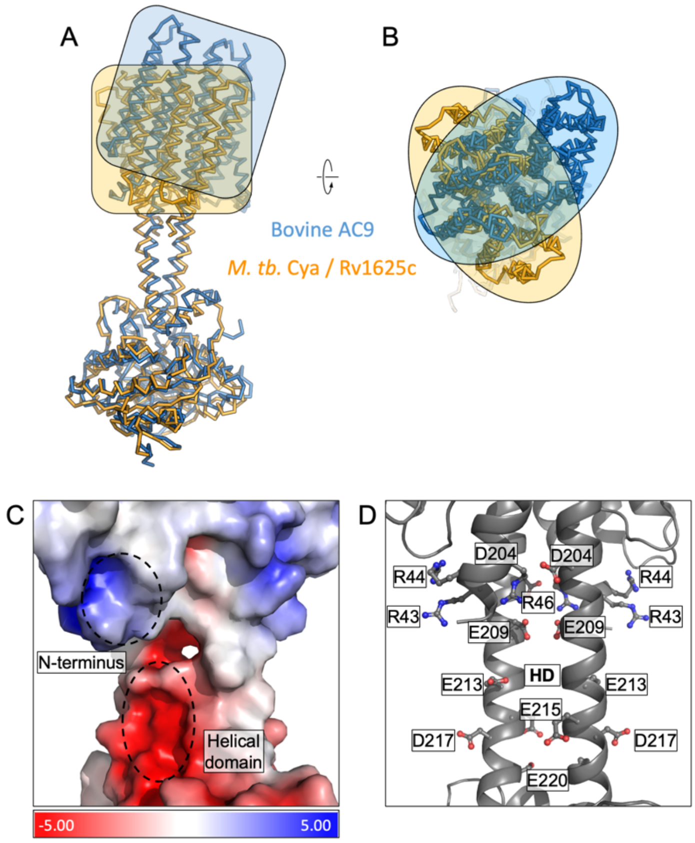


**Figure S5. Comparison of *M. tuberculosis* Cya and bovine AC9. (A)** Structural alignment of Cya and bovine AC9 using their catalytic domains. The TM regions show considerable deviation in alignment, highlighted by orange/blue boxes (*left*) and ovals (*right*) outlining the protein shapes. The top view of the alignment shows an approx. 90˚ rotation of the bAC9 TM domain compared to that of Rv1625c further highlighting the global structural difference between the two cyclases. **(A-B)** Vacuum electrostatic charge distribution of the N-terminus and the HD of Cya **(C)**. The positively charge residues of the N-terminus may interact with the negatively charged residues of the HD **(C-D)**.

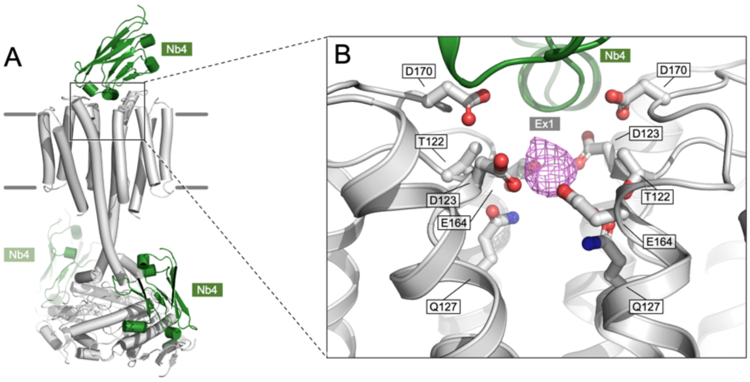


**Figure S6. Density present in the Ex1 site. (A)** An overview of the Cya model in c1 symmetry, indicating the positions of the bound Nb4 chains in green. (**B**) A corresponding zoomed-in view of the site Ex1 in the c1 model, indicating the density feature (magenta) occupying the site in the postprocessed Cya map (c1 symmetry); the corresponding feature in the c2 symmetry map is shown in Fig. 4E. The side chains of the polar residues in the vicinity of the site are indicated as sticks.

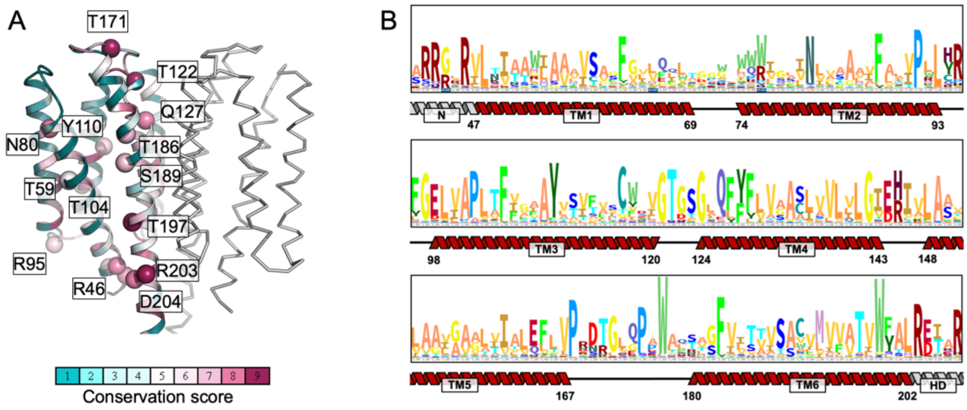


**Figure S7. Sequence conservation of the transmembrane domain residues of mycobacterial Cya homologues. (A)** The conservation scores were calculated using ConSurf *(56)*, using multiple sequence alignment including 170 homologues of Cya from mycobacteria; the sequences of the homologues were obtained by a Blast search. **(B)** The HMM logo of the aligned sequences used in the ConSurf analysis. The secondary structure features indicate the relative positions of the residues in the TM domain.

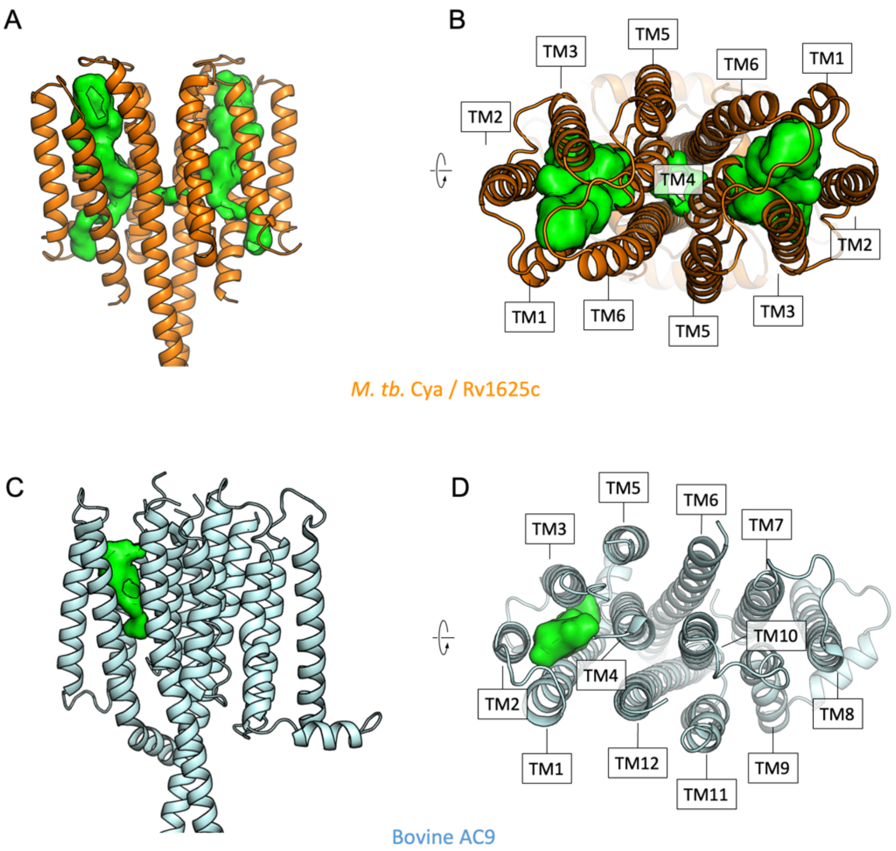


**Figure S8. Comparison of the intramembrane cavities formed by Cya and AC9. (A)** A side view of Cya (light orange) with the Ex2 cavity (bright green) depicted with a cavity detection of 2.5 Å solvent radius and a cavity detection cutoff of 4 Å solvent radius (*left*). The cavity is discontinuous due to stearic constriction, yet prominent and could potentially accommodate a sufficiently small molecule. (*right*). **(B)** A view of the Ex2 cavity perpendicular to the plane of the membrane. The cavity is formed by TM helices 1, 2, 3, 4 and 6. **(C-D)** Same as A-B, for bovine AC9 (light blue) The cavity in bAC9 is less prominent but located in a similar position. However, any potential structural changes could cause the cavity to open (*right*). The cavity in bAC9 is formed by TM helix 1, 2, 3 and 4 (D).

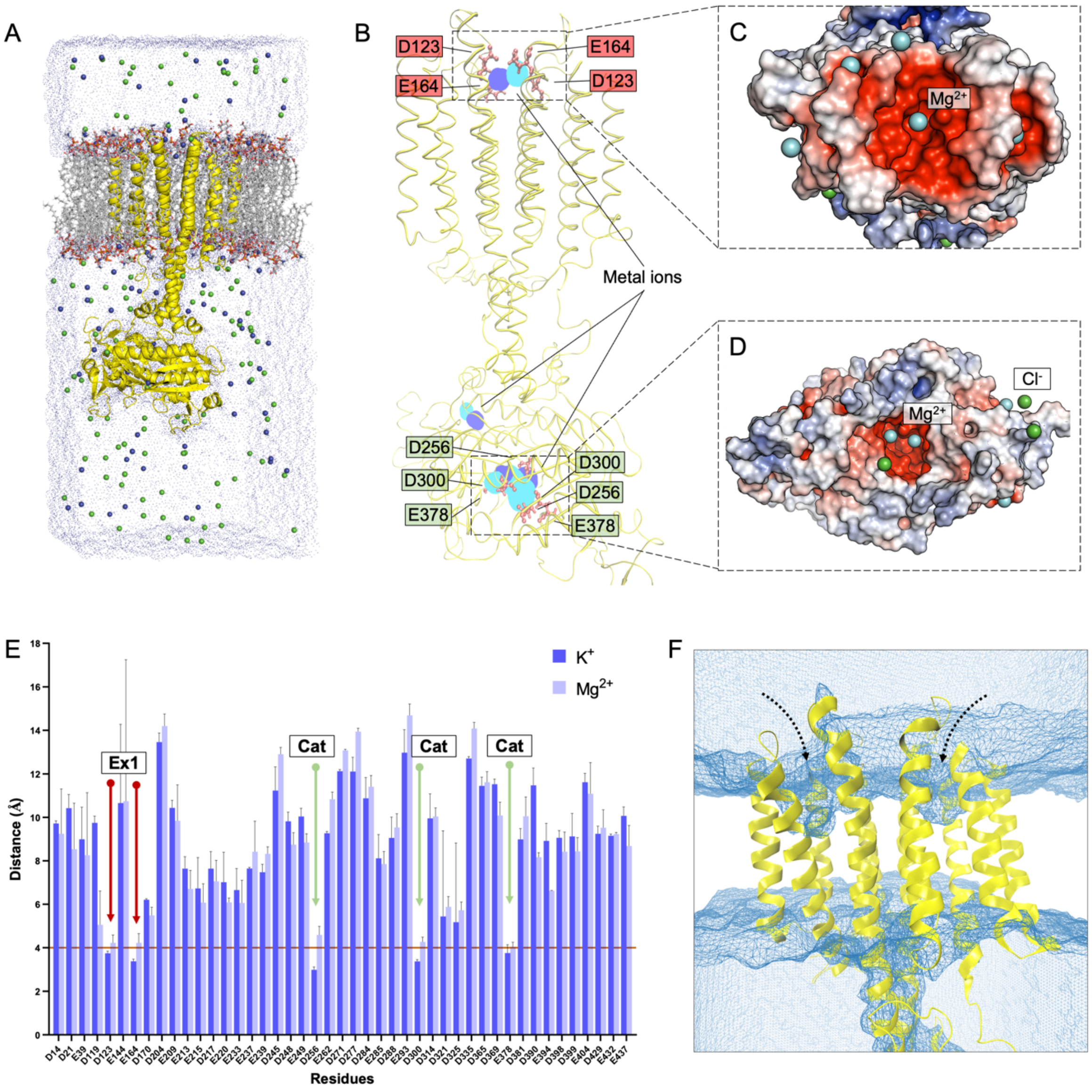


**Figure S9**. **Molecular dynamics simulations of Cya. (A)** Illustration of Cya embedded in lipid bilayer, solvated with TIP3 water molecules and neutralized with 0.15 M KCl or $MgCl_2$ for molecular dynamics (MD) simulations. **(B)** Overlay of potassium (blue) and magnesium (cyan) occupancy maps from independent simulations fitter on the reference structure (yellow). **(C, D)** Depiction of $Mg^{2+}$ and $Cl^-$ ions present within 4 Å of Ex1 site and of the catalytic pocket of Cya at the end of a 200 ns simulation. **(E)** Average distances between negatively charged residues and $K^+$ (light blue) or $Mg^{2+}$ ions (cyan) during the course of a simulation. **(F)** The density grid of water molecules (blue mesh) showing the partial solvent accessibility of Ex2 pocket during simulation.

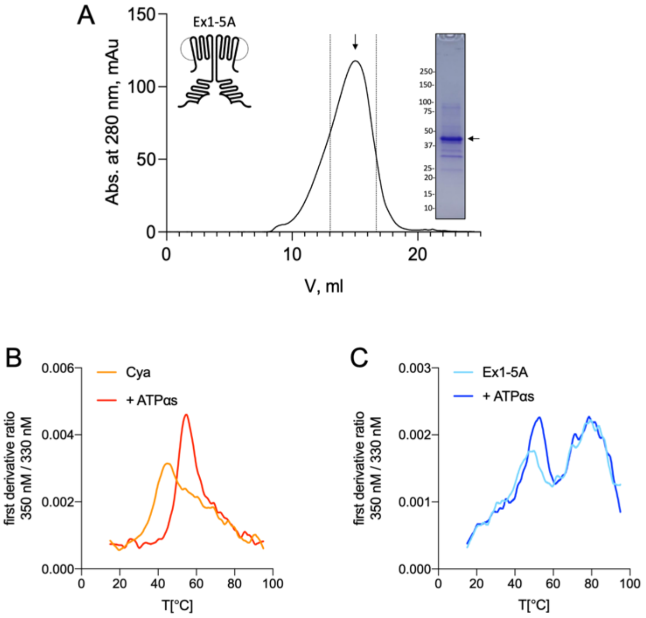


**Figure S10. Purification and stability of Cya mutant Ex1-5A. (A)** Size exclusion chromatography (SEC) and SDS PAGE of Ex1-5A. The dashed lines indicate the selected fractions of the purified protein sample. **(B)** Analysis of protein thermostability, performed using Prometheus Panta instrument, shows that wild-type Cya has an apparent melting temperature of ~42˚C, and is stabilized by addition of the nucleotide analogue (ATPαS), resulting in the shift of the first derivative ratio (350 nm / 330 nm) peak towards higher temperatures. Here, a representative experiment is shown (the average $T_m$ values are shown in Fig. 5). **(C)** Same as in B, for Ex1-5A mutant. The mutant displays a different melting profile, with additional peaks appearing in the high temperature range. Addition of ATPαS shifts the first peak towards high temperatures.

**Table S1.** Cryo-EM analysis and statistics

| Instrument | FEI Titan Krios / Gatan K3 / GIF Quantum LS | |
|---|---|---|
| Magnification | 130000x | |
| Voltage (kV) | 300 | |
| Electron Dose (e-/Å2) | | |
| Dataset 1 | 54 e-/Å$^2$ | |
| Dataset 2 | 47 e-/Å$^2$ | |
| Dataset 3 | 44 e-/Å$^2$ | |
| Defocus range (µm) | -0.5 to -3.0 | |
| Pixel size (Å) | 0.66 | |
| Refinement | | |
| Number of particles | 646042 | |
| Map symmtery | **C2** | **C1** |
| Map FSC, 0.143 | 3.57 | 3.83 |
| Map sharpening b-factor (Å) | -152.693 | -155.051 |
| Model to map FSC, 0.5 | 3.9 | 4.12 |
| Map CC (mask) | 0.69 | 0.66 |
| Model composition, atoms (hydrogen atoms) | 15182 (7560) | 16928 (8413) |
| Protein residues/ligands | 990/6 | 1107/6 |
| Bond length, R.M.S.D. | 0.003 | 0.009 |
| Bond angle, R.M.S.D. | 0.642 | 1.656 |
| Validation | | |
| MolProbity score | 1.46 | 3.06 |
| Clash score | 8.5 | 23.86 |
| Rotamer outliers (%) | 0.77 | 11.24 |
| Mean B-factors protein / ligand | 46.29 / 39.85 | 83.33 / 77.95 |
| Ramachandran plot | | |
| Favoured (%) | 98.16 | 94.24 |
| Allowed (%) | 1.23 | 4.94 |
| Outliers (%) | 0.61 | 0.82 |

**Table S2.** X-ray data analysis and statistics

| | Rv1625c(sol)-NB4 |
|---|---|
| Wavelength | 0.999879 |
| Resolution range | 47.4 - 1.973 (2.044 - 1.973) |
| Space group | P 65 2 2 |
| Unit cell | 94.793 94.793 119.655 90 90 120 |
| Total reflections | 774280 (34068) |
| Unique reflections | 21581 (1491) |
| Multiplicity | 35.9 (22.8) |
| Completeness (%) | 93.96 (66.85) |
| Mean I/sigma (I) | 25.4 (1.5) |
| Wilson B-factor | 44.58 |
| R-merge | 0.09 (1.587) |
| CC1/2 | 1.0 (0.685) |
| Reflections used in refinement | 21583 (1491) |
| Reflections used for R-free | 1077 (75) |
| R-work | 0.2056 (0.3127) |
| R-free | 0.2443 (0.3727) |
| Number of non-hydrogen atoms | 2537 |
| Macromolecules | 2325 |
| Ligands | 57 |
| Solvent | 155 |
| Protein residues | 296 |
| RMS(bonds) | 0.005 |
| RMS(angles) | 0.80 |
| Ramachandran favored (%) | 98.62 |
| Ramachandran allowed (%) | 1.03 |
| Ramachandran outliers (%) | 0.34 |
| Rotamer outliers (%) | 2.85 |
| Clashscore | 8.10 |
| Average B-factor | 58.38 |
| Macromolecules | 57.77 |
| Ligands | 63.98 |
| Solvent | 65.44 |
